# Shared genetic aetiology between cognitive performance and brain activations in language and math tasks

**DOI:** 10.1101/386805

**Authors:** Yann Le Guen, Marie Amalric, Philippe Pinel, Christophe Pallier, Vincent Frouin

## Abstract

Cognitive performance is highly heritable. However, little is known about common genetic influences on cognitive ability and brain activation when engaged in a cognitive task. The Human Connectome Project (HCP) offers a unique opportunity to study this shared genetic etiology with an extended pedigree of 785 individuals. To investigate this common genetic origin, we took advantage of the HCP dataset, which includes both language and mathematics activation tasks. Using the HCP multimodal parcellation, we identified areals in which inter-individual functional MRI (fMRI) activation variance was significantly explained by genetics. Then, we performed bivariate genetic analyses between the neural activations and behavioral scores, corresponding to the fMRI task accuracies, fluid intelligence, working memory and language performance. We observed that several parts of the language network along the superior temporal sulcus, as well as the angular gyrus belonging to the math processing network, are significantly genetically correlated with these indicators of cognitive performance. This shared genetic etiology provides insights into the brain areas where the human-specific genetic repertoire is expressed. Studying the association of polygenic risk scores, using variants associated with human cognitive ability and brain activation, would provide an opportunity to better understand where these variants are influential.

## Introduction

Language and math functions in humans are extensively studied in fundamental neuroscience as distinctive abilities of human lineage. They are frequently assessed through neuroimaging to provide endophenotypes^1,2^. They are used as a way to classify the broad behavioral symptoms of language impairments into stable phenotypes that in turn are candidates to search for potential associations with either medical treatment responses or genetic profiles^3,4^. Structural properties observed using magnetic resonance imaging (MRI) or activations observed with functional MRI (fMRI) have been used to produce such endophenotypes^5^. These can reveal differences between control and disease groups in language-specific regions^6^ or distinguish disorder subtypes such as grammatical-SLI (specific language impairment)^7^.

Imaging-genetics resources, such as the Human Connectome Project (HCP) provide an unprecedented opportunity to study the variability of such endophenotypes in control subjects, as well as to determine their potential heritability or association with genetics. Following up on these ideas, we first proposed to study the additive genetic variance involved in fMRI activation differences among typically developed individuals. We used the pedigree data from the HCP language comprehension and verbal math fMRI tasks. These tasks recruit regions directly implicated in brain disorders, such as Broca’s area in SLI^8^, the angular gyrus in developmental dyslexia^1^ and the intraparietal in dyscalculia^2^. A few studies have already attempted to estimate the narrow sense heritability of brain activations for various tasks. They notably include digit and n-back working memory^9,10^, visual math subtraction^11^, and stimuli such as written words, faces and spoken language^12^. However, these studies had relatively small sample sizes for reliably detecting heritability estimates ranging between 25 and 50%. The previous samples included 30 subjects (10 triplets of male monozygotic (MZ) twins with one additional brother)^9^, 64 subjects (19 MZ and 13 dizygotic (DZ) pairs)^11,12^ or 319 subjects (75 MZ and 66 DZ pairs, 37 unpaired)^10^. In addition to including a larger sample size, the HCP data were processed using state of the art methods, providing 2-mm isotropic resolution and finer inter-individual registration. In particular, the so-called grayordinate activations are computed on the surface of the cortex^13^ for each individual, and inter-subject fMRI alignment is performed using areal-feature-based registration^14^. The grayordinate approach refers to fMRI analyses performed on the cortical surface, as opposed to a volume-based approach. The same idea was applied to build the multimodal parcellation of the human cerebral cortex^15^ onto which we decomposed our heritability analyses, enabling us to map the genetic influence on fMRI activations on a very fine scale.

Furthermore, it is known that neural activation endophenotypes from MRI may reflect not only impairments in language but also differences in cognitive scores. For example, weaker left-lateralizations have been reported for some developmental language disorders^16^, and in normal populations, fMRI activations during simple tasks correlate with various cognitive scores. Notably, single digit calculation fMRI activations are predictive of high school math scores^17^, and the fronto-parietal functional connectivity in children performing a task that required them to match Arabic numbers to an array of dots correlated with their score on a standardized math test^18^. Using HCP data and in line with these approaches, we show in this paper how variations in language related fMRI activations correlate with cognitive abilities assessed by the median reaction time (RT), average accuracy and difficulty level during the HCP language and math tasks. Remarkably, recent studies have shown that, beyond the age-related heritability of general cognitive ability^19,20^ and of various indicators of academic performance^21^, these scores are highly pleiotropic^19,22^ [pleiotropy occurs when one gene regulates one or more phenotypic traits].

This raises the question of the potential pleiotropy between neural activations and cognitive abilities. Thus, as a second contribution, we studied the shared genetic variance of fMRI activations and cognitive performance scores measured during the MRI session or behavioral scores acquired independently from the task. We studied behavioral variables measured by the HCP using standardized tests from the National Institute of Health (NIH): fluid intelligence, working memory, and language assessments such as vocabulary comprehension and oral reading decoding. Details of these variables and their heritability estimates can be found in Table S1, and how well they correlate phenotypically and genetically with the behavioral scores measured during the task is reported in Table S2.

Recent genome wide association studies have unveiled new loci and genes influencing human cognitive performance (e.g. human intelligence^23^, general cognitive function^24,25^ and educational attainment^25^) and possibly intelligence as a construct in differential psychology^19^. However, for these human-specific characteristics, little is known about the underlying integration mechanism of molecular functions or the brain areas where they are most influential. The shared genetic etiology investigated in this work provides new perspectives to decipher the basis of cognitive abilities such as language in humans. This study had two major aims: (1) to estimate the heritability of fMRI activations during story comprehension and math tasks; and (2) to determine the shared genetic etiology between these activations and cognitive performance.

## Results

### Task fMRI Activations in MATH and STORY tasks

Figure 1 shows the activations for MATH (vs the intercept of the general linear model (GLM) being considered as baseline), STORY and the contrast STORY - MATH. The intercept reflects the mean of the residual BOLD time series after removing variance explained by all other regressors. Both tasks show clear activations in the planum temporale and Heschl’s gyrus area, reflecting the fact that the stimuli were presented in the auditory modality. The MATH task, in which participants were requested to perform addition and subtraction, activates areas traditionally implicated in mathematical calculations, that is, the intraparietal sulcus, the middle frontal and the inferior temporal regions^26,27^. The story listening task activates the language understanding network, encompassing bilateral temporal regions and left frontal regions^28,29^. As expected, the group activations for the STORY task are more left lateralized, notably in the left posterior superior temporal and inferior frontal regions, which correspond to Wernicke’s and Broca’s areas, respectively. Moreover, regions implicated in inhibition networks are also activated by these tasks^30,31^, notably the middle frontal gyrus in the math task and the medial prefrontal cortex, implicated in motivation and execution, and above the anterior cingulate cortex, controlling selective attention^32,33^. In addition, both tasks activate complementary networks; in particular, the math task deactivates the semantic and episodic memory processes, known as the default mode network, which is also active in resting or passive states^34^. This last remark makes the STORY-MATH contrast particularly relevant for studying the genetic influence on activation specifically elicited by math and story tasks.

**Figure 1.**
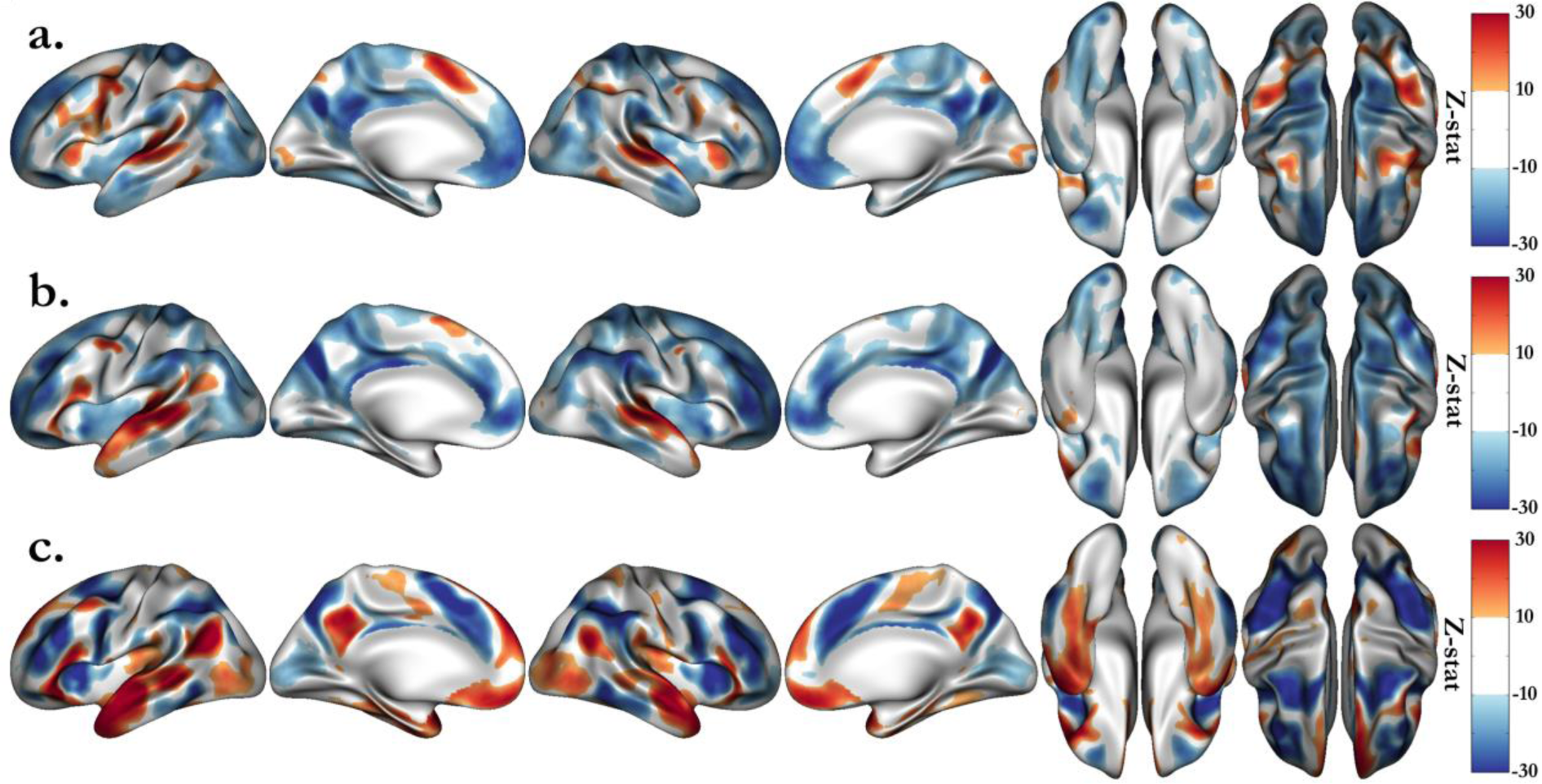
Group average activations for the HCP language tasks, including MATH (a.) and STORY (b.) blocks, and the STORY-MATH contrast (c.). Group maps are shown with a lower threshold of z = ± 10 and saturation from z = 30 to introduce the main areas activated by the tasks. Due to the large number of subjects, the associated p-values are significant; we arbitrarily set the thresholds to emphasize the regions that are known to be recruited by these tasks.

### Univariate Genetic Analyses

We performed a cortex-wise heritability analysis on the median activation (β-value) in the 360 areals of the HCP multi-modal parcellation. After stringent Bonferroni correction (p < 0.05/360 ≈ 1.4 ≈ 10^−4^), we found 54 regions whose activations during the MATH task are heritable and 46 regions for the STORY task. These results are summarized in Figure 2 and heritability estimates are included in Tables S3-S6. The details and names of the areals can be found in the Supplementary Information of the paper describing the multimodal parcellation of the human cerebral cortex^15^. For the MATH (resp. STORY) task, the heritability estimates range from 0.23 to 0.45, with the maximum in the left “Area PGp” corresponding to the angular gyrus (resp. 0.22 to 0.55, with the maximum in the left “PeriSylvian Language Area”). In addition, we performed heritability analysis using the median z-stat value in each areal instead of the median parameter estimate (β-value) and obtained similar results (Fig S1).

**Figure 2.**
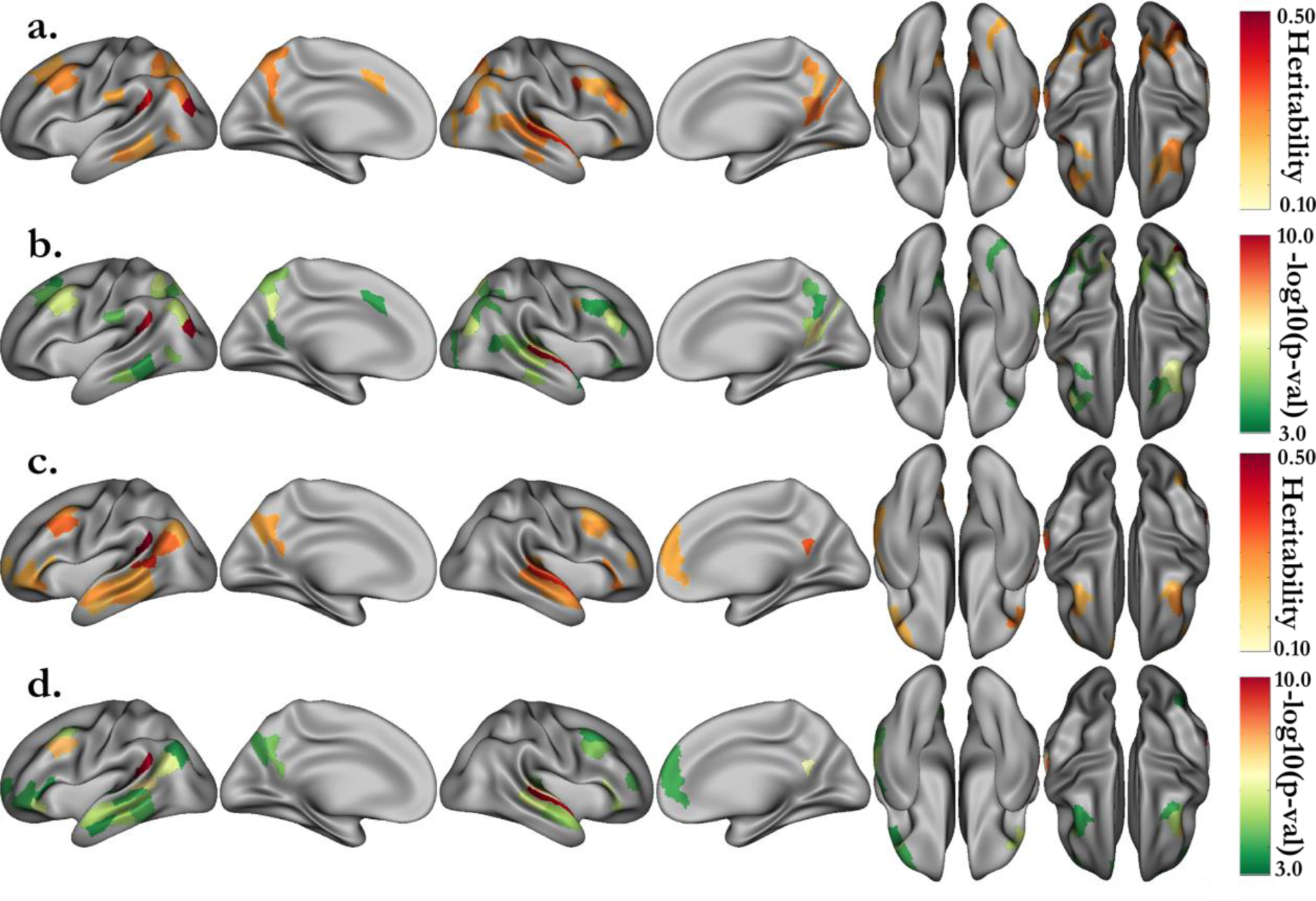
Heritability estimates for the activations of the MATH (a.) and STORY (c.) tasks, and their associated p-values (respectively, b. and d.). Only the estimates significant after correction (p < 0.05/360, with 180 areals in each hemisphere) are displayed. Activations correspond to the median parameter estimate (β) in each areal of the HCP multimodal parcellation.

The univariate genetic analysis of the activations associated with verbal math emphasizes mainly areals spanning the math network, including the intraparietal sulcus, middle frontal, inferior temporal and angular gyri. The analysis of activations associated with story comprehension distinctively underlines regions of the language network as bilaterally heritable. Among these regions are the superior temporal sulcus dorsal and ventral parts, Brodmann area (BA) 47 in Broca’s area, and the middle frontal gyrus at the junction with the precentral sulcus. Interestingly, the heritability networks of the MATH and STORY tasks overlap very little except in the auditory cortex, around the planum temporale, in the frontal cortex (BA 8), and in the inferior temporal region.

Table 1 presents the heritability estimates of the behavioral scores gathered during the MRI scans. The global accuracy on the HCP language tasks, averaging the scores in the MATH and STORY tasks, was significantly heritable, with h² = 0.34, close to the traditionally high heritability estimate of cognitive performance^35,36^. The heritability estimates for the median reaction time (RT) were approximately 0.2. Furthermore, RT Story and RT Math were significantly correlated (phenotypic correlation: ρ_p_ = 0.34, genetic correlation: ρ_g_ = 0.45, Table S2). Regarding the accuracy and average difficulty level of the HCP language task, we observed that the MATH task variables have higher heritability estimates than those of the STORY task. This result might indicate a higher genetic influence on performance during simple arithmetic tasks than during language comprehension. However, this result needs to be considered in light of the different distribution patterns of MATH and STORY accuracies. The STORY accuracy reported by the HCP displays discrete values and might not be sufficiently informative (Fig S2). Table S2 underlines a significant correlation between math and story accuracies (ρ_p_ = 0.15, ρ_g_ = 0.27). The discrete distribution of STORY accuracy likely occurs because each story lasted approximately 20-30s, few story questions were presented to the subjects, and most subjects tended to choose the correct answer in the two-alternative forced-choice question.

**Table 1.** Heritability estimates for the behavioral scores associated with the tasks. Language accuracy and reaction time (RT) correspond to the average of the respective MATH and STORY variables. The p-values associated with the covariates related to age and sex, ethnic group and education level are also displayed.

| Trait | h<sup>2</sup>±SE (p) | Age | Age<sup>2</sup> | Sex | Age*Sex | Age<sup>2</sup>*Sex | Hispanic | Educ | h<sup>2</sup>cov% |
| --- | --- | --- | --- | --- | --- | --- | --- | --- | --- |
|  |  | p-val |  |  |  |  |  |  |  |
| Language Accuracy | 0.34±0.06 (2.3·10<sup>-8</sup>) | 0.49 | 0.75 | 0.01 | 0.96 | 0.22 | 0.97 | 1.2·10<sup>-7</sup> | 7.2 |
| Language RT | 0.22±0.07 (8.4·10<sup>-4</sup>) | 0.44 | 0.91 | 0.69 | 0.85 | 0.95 | 0.71 | 0.04 | 0.5 |
| Math Accuracy | 0.4±0.06 (1.6·10<sup>-10</sup>) | 0.5 | 0.96 | 9.2·10<sup>-4</sup> | 0.81 | 0.39 | 0.76 | 1.9·10<sup>-6</sup> | 7.2 |
| Math Difficulty Level | 0.33±0.07 (1.0·10<sup>-6</sup>) | 0.53 | 0.17 | 0.02 | 0.61 | 0.05 | 0.42 | 8.7·10<sup>-8</sup> | 7.9 |
| Math Median RT | 0.17±0.07 (7.4·10<sup>-3</sup>) | 0.6 | 0.41 | 0.78 | 0.82 | 0.69 | 0.26 | 0.01 | 1.1 |
| Story Accuracy | 0.18±0.06 (1.6·10<sup>-3</sup>) | 0.78 | 0.48 | 0.86 | 0.76 | 0.23 | 0.61 | 2.2·10<sup>-3</sup> | 1.4 |
| Story Difficulty Level | 0.33±0.07 (1.3·10<sup>-6</sup>) | 0.66 | 0.43 | 0.81 | 0.44 | 0.28 | 0.91 | 0.11 | 0.0 |
| Story Median RT | 0.2±0.07 (1.4·10<sup>-3</sup>) | 0.41 | 0.63 | 0.39 | 0.59 | 0.68 | 0.45 | 0.39 | 0.0 |

### Bivariate Genetic Analyses

We performed bivariate genetic analyses to quantify the shared genetic influence between intellectual performance, represented by the behavioral measures, and the neural activation in each areal. The genetic correlation estimates are usually subject to substantial sampling errors and therefore inaccurate. The large sample size of the HCP offers the opportunity to reduce the standard errors. The distribution of STORY accuracy values is concentrated on a small number of values (Fig S2), thus, we chose to use the average of the STORY and MATH accuracies as the behavioral score to characterize the individual performance. Thus, the results presented here concern the relationship between this average score and the activations or deactivations revealed by the contrast STORY-MATH (activation map shown in Figure 1c). As a first step of our analysis and to filter out the areals for which the neural activation was not significantly correlated with the behavioral score, we computed the phenotypic correlation between these two variables in each areal of the HCP multi-modal parcellation. Figure 3 (a, b) summarizes the phenotypic correlation and associated p-values, for areals significant after Bonferroni correction (p < 0.05/360). The language network is clearly encompassed along the left superior temporal sulcus (STS) and Broca’s area, as well as in the anterior part of the right STS. The activations in the angular gyrus (area PGp), supporting the manipulation of numbers in verbal form^37^, were also significantly correlated with the behavioral scores.

**Figure 3.**
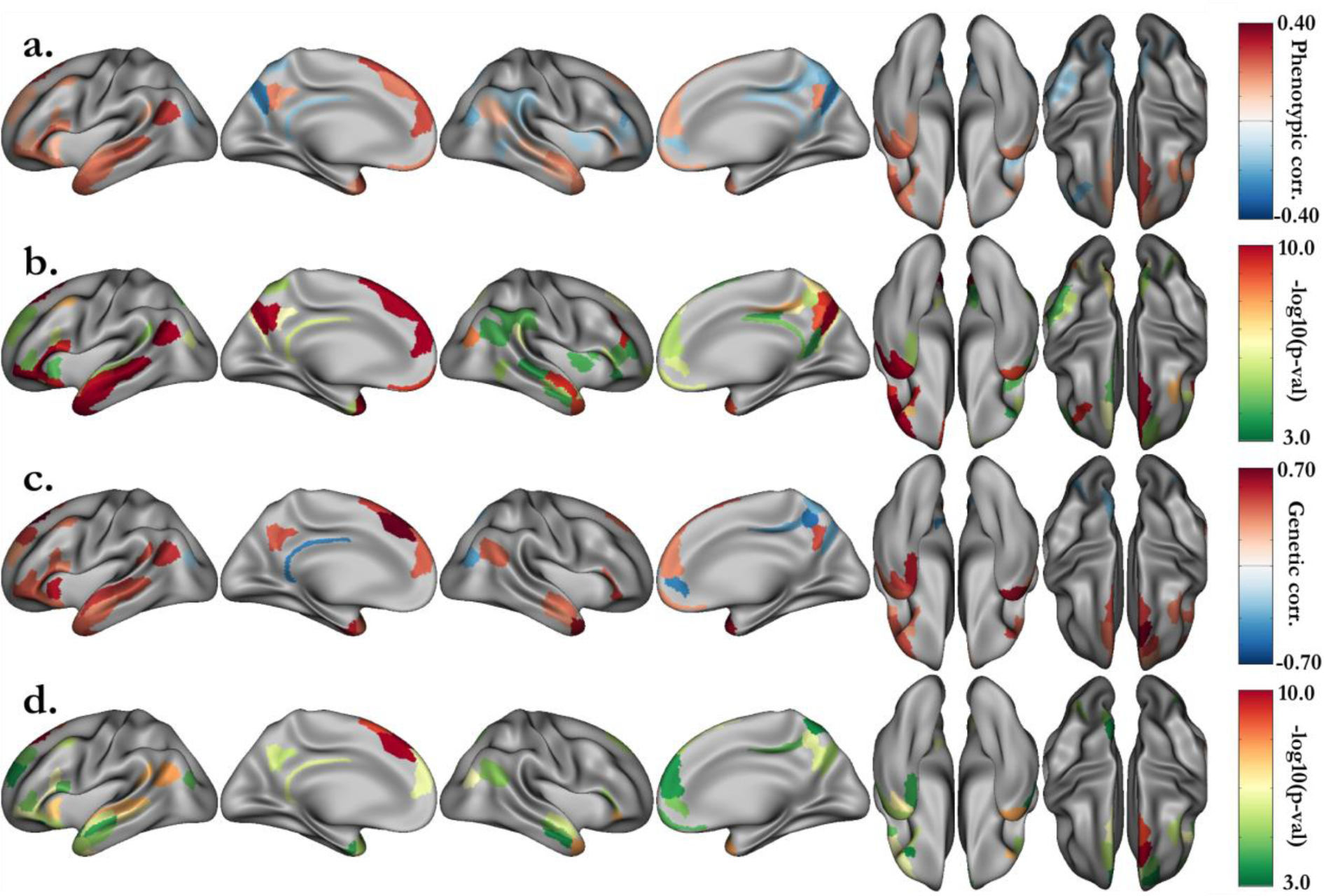
Bivariate genetic analysis results between HCP LANGUAGE task accuracy and activation for the STORY-MATH contrast in each areal. **a.** After strict Bonferroni correction (p < 0.05/360), significant phenotypic correlations between the language task accuracy (average of story and math accuracies) and the median activation of the contrast STORY-MATH in each areal, **b.** with their associated p-values. **c.** Proportion of variability due to shared genetic effects with **d.** their associated uncorrected p-values < 0.05.

Among the 360 areals of the HCP multimodal parcellation, 39 (resp. 38) were significantly phenotypically correlated and were kept for the bivariate analysis in the right (resp. left) hemisphere. The shared genetic variance estimates for these areals are presented Figure 3 (c, d) (p < 0.05 without correction), and detailed values can be found in Tables S7 and S8. With stringent Bonferroni correction the p-value threshold for ρ_g_ is p < 0.05/(39+38) ≈ 6.5 ≈ 10^−4^. Among the areals with significant shared genetic variance, we found the left anterior ventral insular area (AVI, ρ_g=_ 0.61)and the right angular gyrus (PGp, ρ_g=_ −0.40). Noticeably, in the left hemisphere areals, parts of the language network in the posterior STS had activations that shared significant genetic variance with language accuracy. These include the posterior ventral (STSv posterior, ρ_g=_ 0.47) and dorsal (STSd posterior, ρ_g=_ 0.45) parts of the STS, adjacent to the auditory 5 complex area (A5, ρ_g=_ 0.54), the perisylvian language area (PSL, ρ_g_ = 0.47) and the temporo-intraparietal junction (PGi, ρ_g=_ 0.54). On the left hemisphere internal face, we also found the superior frontal language area (SFL, ρ_g=_ 0.61) and, adjacent to this areal the Brodmann 8 decomposed into medial (8Bm, ρ_g=_ 0.75) and lateral (8Bl, ρ_g=_ 0.73) parts. Additionally, we noted two right hemisphere regions implicated in language processing and significantly genetically correlated with the fMRI task average score: the temporal pole (area TG dorsal, TGd, ρ_g=_ 0.65) and the lateral part of Brodmann area 47 (47l, ρ_g=_ 0.51). The latter is adjacent to Brodmann areas 44 and 45 in the inferior frontal, which are connected through the arcuate fasciculus with the language temporal regions.

Additionally, we extended our analysis to behavioral variables measured by the HCP following a standardized NIH protocol. Among these, we selected the variables that are most likely to reflect cognitive performance. Then, we estimated their heritability (Table S1) and correlations with the behavioral scores measured during the fMRI task (Table S2). In this set of variables, fluid intelligence (heritability: h^2^=0.43, correlations with language accuracy: ρ_p_ = 0.36, ρ_g_ = 0.61), working memory (h^2^=0.52, ρ_p_ = 0.34, ρ_g_ = 0.50), vocabulary comprehension (h^2^=0.64, ρ_p_ = 0.40, ρ_g_ = 0.57) and oral reading decoding (h^2^=0.67, ρ_p_ = 0.46, ρ_g_ = 0.67) were the ones with the highest heritability estimates and correlations with the average accuracy of the two fMRI tasks. Thus, we performed a bivariate genetic analysis between the STORY-MATH activations (difference between β_STORY_ and β_MATH_) and these four variables. Regardless of whether one considers the NIH scores or the ones directly related to the fMRI tasks, the study of shared genetic influence with the median activation yields approximately the same set of regions (Figures 4, 5). This observation reinforces our claim that these regions have common genetic roots with the parts of general cognitive performance accounted for by the four cognitive variables under scrutiny, namely fluid intelligence, working memory, vocabulary comprehension and reading decoding.

**Figure 4.**
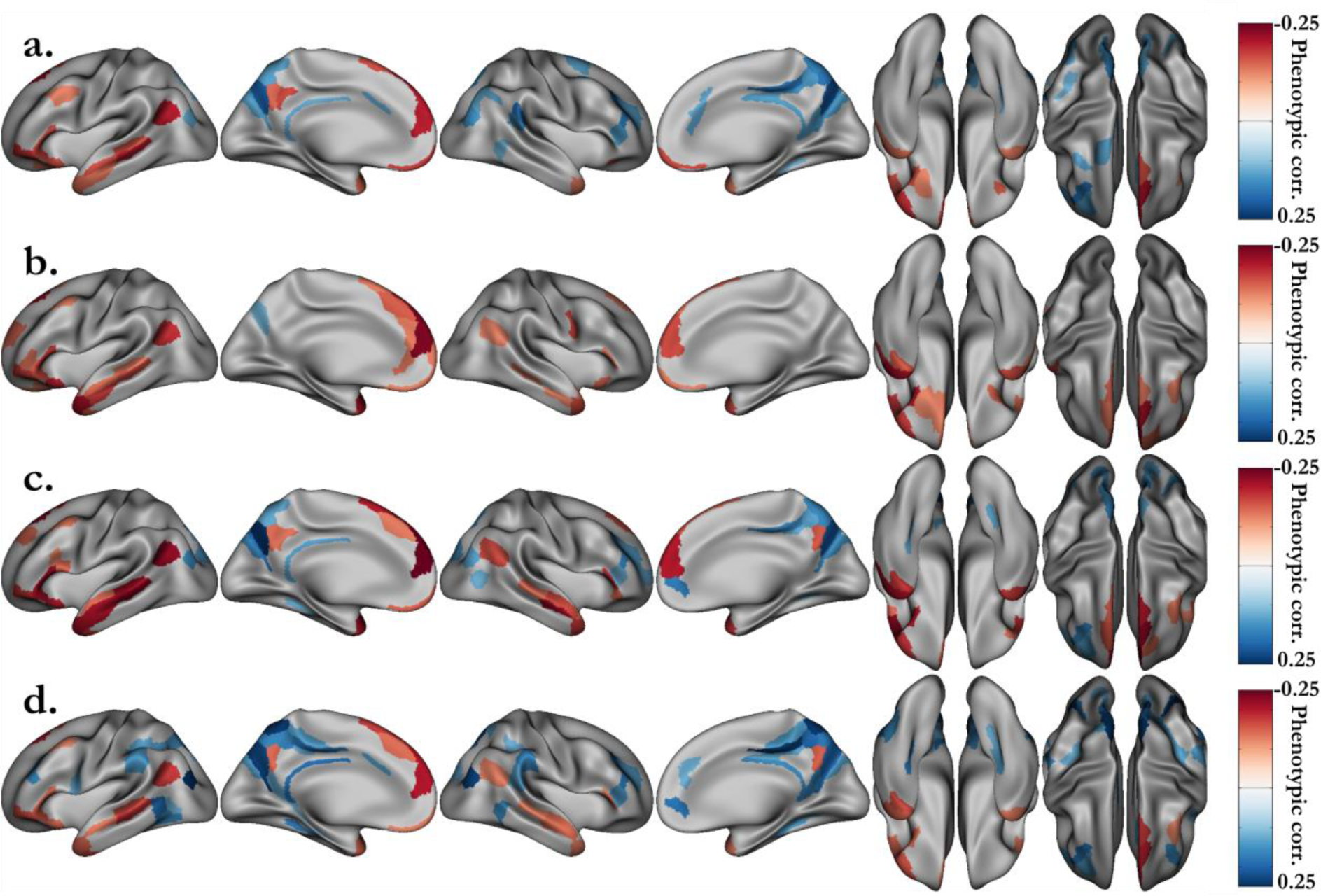
Significant phenotypic correlations between the grayordinate activations of the STORY-MATH contrast and the NIH behavioral scores. **a.** Fluid intelligence (PMAT24_A_CR). **b.** Working memory (ListSort). **c.** Vocabulary comprehension (PicVocab). **c.** Reading decoding (ReadEng). Associated p-values (p < 0.05/360, Bonferroni correction) can be found Fig S3.

**Figure 5.**
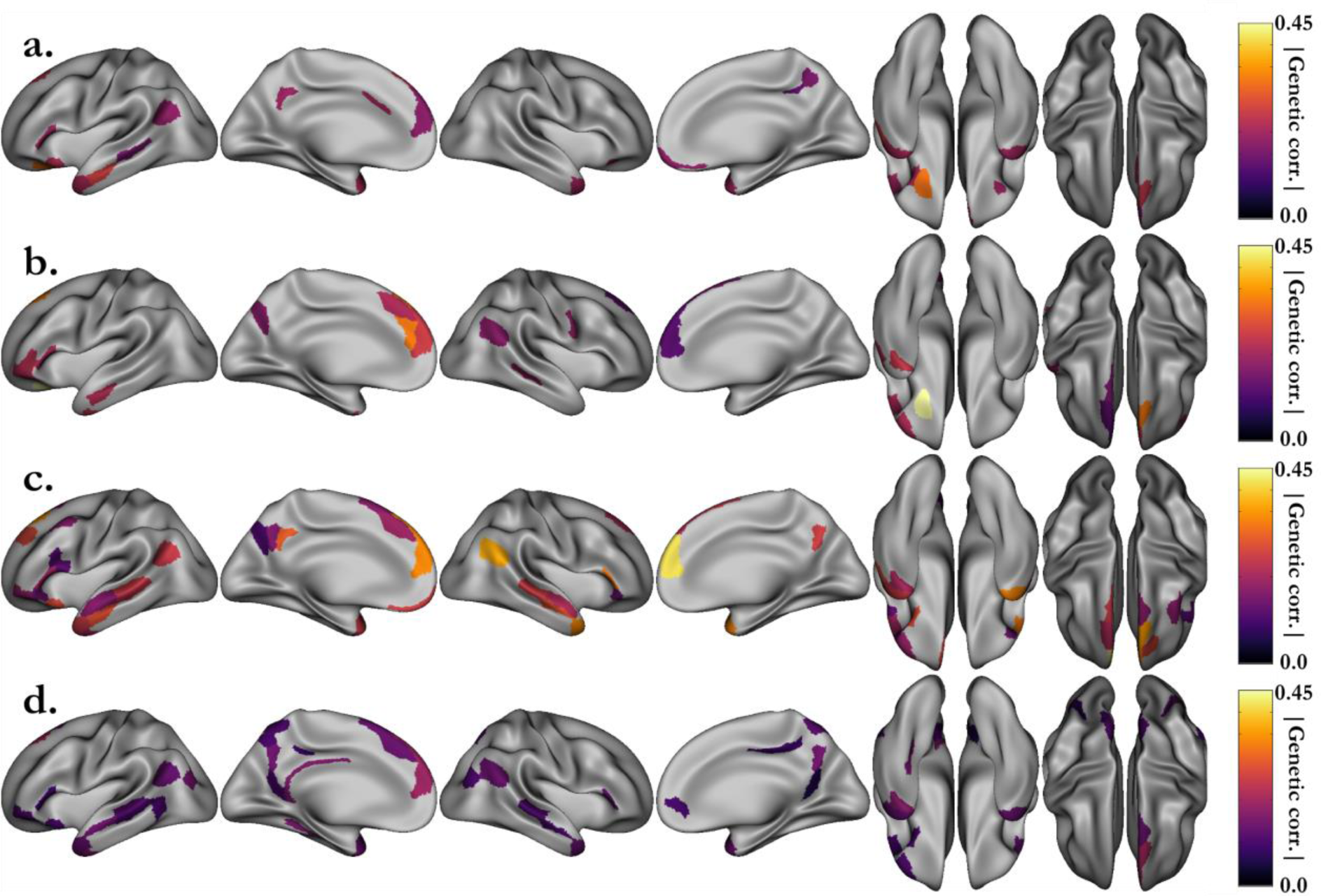
Shared genetic variance (absolute value) between the grayordinate activations of the STORY-MATH contrast and the NIH behavioral scores. **a.** Fluid intelligence (PMAT24_A_CR). **b.** Working memory (ListSort). **c.** Vocabulary comprehension (PicVocab). **c.** Reading decoding (ReadEng). Associated p-values (p < 0.05, uncorrected) can be found in Fig S4. Genetic correlation was investigated only for areals that were significantly phenotypically correlated (Figure 4).

## Discussion

In this paper, we have shown that brain activation pattern in the language and math networks are heritable. Additionally, we highlighted a particular set of regions along the superior temporal sulcus and in the inferior frontal whose activations share a common genetic basis with some aspects of general cognitive ability, assessed through fMRI task accuracy and behavioral scores.

We must emphasize that these results correspond to fMRI activations associated with verbal math and semantic comprehension tasks. Thus, regions not recruited by the tasks cannot be found to be significantly correlated with cognitive ability in our case, because activations in these regions are incoherent across individuals. Notably, the visual word form area, related to literacy, is not activated in our oral tasks because they did not require word reading.

Furthermore, combining data from various cohorts is unfeasible, because neural activations from different tasks are not comparable when estimating inter-individual variance. This highlights the necessity of utilizing large cohorts with standardized fMRI protocols to perform such genetic analyses. To our knowledge, this is the first study to address the heritability of fMRI activation cortex-wise on a multimodal parcellation of the human cerebral cortex. Our results confirm the genetic influence on the formation of neural circuits implicated in language^38^ and math^11^. Using the HCP fine scale parcellation^15^ allowed us, for instance, to distinguish the genetic effects on the temporo-parietal junction implicated in language^28^ (area PGi) and on the adjacent angular gyrus (area PGp), which is particularly involved in the manipulation of numbers in verbal form^37^. Indeed, these two areas, part of BA 39, present different cytoarchitectonic properties, such as a slightly broader layer II for PGi^39^, which might explain their involvement in different tasks. In a previous work, Pinel and Dehaene also found the left angular gyrus and the posterior superior parietal lobule bilaterally to be heritable^11^. Adding to these observations, our results underline a left hemisphere intraparietal specificity, with more heritable areals and slightly higher heritability compared to the right for the MATH contrast. This finding is consistent with results reported by Vogel and colleagues demonstrating a correlation of activations in the left intraparietal sulcus modulated by age, which was not observed in the right intraparietal^40^. Heritability represents the proportion of observed inter-individual phenotypic variance that is explained by genetics. Thus, it might be that inter-individual variance is not sufficiently pronounced in the right hemisphere, whereas activations have evolved over one’s lifetime in the left hemisphere. Overall, the heritability maps for the STORY and MATH tasks pinpoint regions known to be disrupted in neurodevelopmental disorders. For instance, the inferior frontal area and the temporo-parietal junction activations are impaired in developmental dyslexia^1,41^, and the intraparietal region activations are less modulated by the numerical distance between two numbers being compared in developmental dyscalculia^2,42,43^. Highlighted areas might provide new insights into brain regions where normal gene expression might be disrupted, leading to brain dysfunction and neurodevelopmental disorders. Frequently replicated genes associated with neurobehavioral disorders, such as developmental dyslexia or SLI, likely play such a role in structural brain maturation by interfering with neuronal migration and neurite growth^4^.

Several studies have already described some phenotypic correlations between cognitive abilities and neural activations in language^44–46^ and math^17,18,47,48^. Our study replicates these observations, notably the correlation with language processing regions, including Broca’s area and the posterior superior temporal gyrus^44^. Moreover, we estimated the genetic proportion in these phenotypic correlations. Hence, we demonstrated a shared genetic etiology between brain activations and cognitive performance, assessed in our study by the following tests: fluid intelligence, working memory, vocabulary comprehension and reading decoding. Interestingly, in the right hemisphere, mainly the anterior STS was found to be genetically correlated with the language task accuracy. This result seems consistent with the hypothesized role of the right anterior STS in the processing of prosody or figurative language, likely involved in the Aesop’s fable metaphors presented to the subjects^49^.

The observed genetic correlations shed light on the genetic links between cognitive performance and activation level in cognitive task-related fMRI. These links might be related to the development and maturation of myelin, enhancing brain connectivity. In children with difficulties processing syntactically complex sentences, arcuate fasciculus maturation was incomplete compared to adults^50,51^. Thus, we could look for additive genetic effects implicated in the various levels of fiber tract maturation, which improves brain connectivity and efficiency. Indeed, Skeide and colleagues reported an example of such a genetic risk variant for dyslexia. They showed that this variant is related to the functional connectivity of left fronto-temporal phonological processing areas during the resting state^52^. Similarly, children with higher arithmetic scores present a more mature response modulation in their left intraparietal lobe^47^. Our study suggests that a proportion of the observed inter-individual variance in cognitive performance partly results from the same additive genetic effects as those contributing to brain activation variance. The moderate shared genetic basis suggests that a crucial interaction occurs between the environment and gene networks to enable the brain to develop to its full potential.

Recently, with the emergence of large cohorts, such as UKBiobank, new loci and genes influencing human cognitive ability have been discovered^23–25^. However, little is known about how these genes contribute to this human-specific trait. Our study pinpoints brain regions where activations genetically correlate with global cognition scores. These regions might help elucidate the mechanism in which these genes are implicated. When the HCP genotyping data are released, a polygenic score of these newly discovered variants could be used to determine the explained proportion of the neural activation variance in these regions.

## Material and Methods

### Subjects

This study utilized the dataset of the Human Connectome Project (HCP). The HCP scans and data were released in April 2017 (humanconnectome.org). The details of the release are available in the HCP reference manual. In this project, 1046 subjects aged between 22 and 37 years old (µ ± σ = 28.8 ± 3.7 years) completed the fMRI language task in the HCP S1200 release. To avoid population stratification, we only included the 785 Caucasian individuals (372/413 M/F) that were classified as race=“White” by the HCP; among these, the ethnicity of 69 was “Hispanic/Latino”. This subgroup of the HCP contains 178 twin pairs (117 monozygotic twins (MZ) with 103 siblings and 61 dizygotic twins (DZ) with 61 siblings and 1 half sibling), 203 siblings, 1 half sibling and 60 unpaired individuals. The unpaired individuals did not contribute to the genetic parameter estimation but allowed a more accurate estimation of mean and variance effects. Subjects were chosen by the HCP consortium to represent healthy adults beyond the age of major neurodevelopmental changes and before the onset of neurodegenerative changes ^53^. They underwent a battery of tests to determine if they met the inclusion/exclusion criteria of the HCP ^53^. All subjects provided written informed consent on forms approved by the Institutional Review Board of Washington University. All of the following methods were carried out in accordance with relevant guidelines and regulations. All experimental MRI protocols were approved by the Institutional Review Board of Washington University.

#### Data availability statement

HCP (https://db.humanconnectome.org) is a publicly available dataset. Investigators need to apply to be granted access to restricted data.

### Statistical power analysis

The statistical power of an extended pedigree study should not be confused with that of a cohort of unrelated subjects. In the latter case, the software often used is GCTA (genome wide complex trait analysis)^54^, which enables one to easily compute the statistical power^55^. In an extended pedigree, the the statistical power computation is a hard problem because non-independence among relatives must be taken into account^56^. To the best of our knowledge, there is no consensus method to compute the *a priori* statistical power for studies on imaging genetics in the case of a large pedigree cohort. For the sake of comparison, we propose to compare our study with previously published studies. Previous heritability studies that used approximately 800 subjects from the HCP, roughly as much as in our sample, obtained standard errors between 0.06 and 0.08 for phenotypes with heritability between 0.20 and 0.45^57,58^. Using the GCTA Power Calculator^55^ with the default value for the variance of the single nucleotide polymorphism (SNP) derived genetic relationship matrix of 10^−5^, this range of standard errors and heritability estimates requires a sample size of 6000-7000 unrelated individuals. To allow comparison with other pedigrees studies, we computed a relatedness summary parameter (RSP) using the pedRSP R package^59^. We determined that our sample of 785 individuals is equivalent to an effective number of 1,922 pairs of siblings (RSP of our sample: V_λ_ = 0.649, V_q_ = 0.671, γ = 0.73; see 59 for details of the parameter’s meaning).

Additionally, we want to emphasize that modeling non-additive genetic effects^60^ is not possible when genetic data are not available. In a pedigree study such as the one currently available from the HCP, the only information available is the relationship between individuals.

### Image acquisition and processing

MR images were acquired using a 3T Connectome Scanner, adapted from Siemens Skyra, housed at Washington University in St Louis, using a 32-channel head coil. T1-weighted images with 256 slices per slab were acquired with the three-dimensional magnetization-prepared rapid gradient echo (3D-MPRAGE) sequence: TR=2400 ms, TE=2.14 ms, TI=1000 ms, flip angle=8°, FOV=224×224 mm, and resolution 0.7 mm isotropic. T2-weighted images, 256 slices per slab, were acquired with a 3D-T2SPACE sequence: TR=3200 ms, TE=565 ms, variable flip angle, FOV=224×224 mm, and resolution 0.7 mm isotropic. fMRI data acquisition parameters were as follows: TR=720 ms, TE=33.1 ms, flip angle=52 deg, BW=2290 Hz/Px, in-plane FOV=208×180 mm, 72 slices, and 2.0 mm isotropic voxels, with a multi-band acceleration factor of 8. Two runs of each task were acquired, one with right-to-left and the other with left-to-right phase encoding 2; each run interleaved 4 blocks of a story task with 4 blocks of a math task. The lengths of the blocks varied (average of approximately 30 seconds), but the task was designed so that the math task blocks matched the length of the story task blocks, with some additional math trials at the end of the task to complete the 3:57 (min:sec) run.

The details of the HCP data analysis pipelines are described elsewhere^13,61^. Briefly, they are primarily built using tools from FSL^62^ and Freesurfer^63^. The HCP fMRIVolume pipeline generates “minimally preprocessed” 4D time series that include gradient unwarping, motion correction, fieldmap-based EPI distortion correction, brain-boundary-based registration of EPI to structural T1-weighted scans, non-linear (FNIRT) registration into MNI152 space, and grand-mean intensity normalization^61^. For the S500 release, two smoothing approaches were chosen by the HCP: volume-based smoothing or smoothing constrained to the cortical surface and subcortical gray-matter parcels. For the former, standard FSL tools can be applied for analysis, while for the latter, the HCP adapted these tools to this the ‘grayordinate’ approach^13,61^. The grayordinate approach refers to fMRI analyses performed on the cortical surfacen, as opposed to a volume-based approach. This is more accurate spatially because activation occurs in gray, not white, matter. Unconstrained volume-based smoothing causes blurring effects by mixing signals from cortex regions adjacent in volume but not on the surface. For these reasons, our study analyses were carried out on the surface of the cortex.

The HCP fMRISurface pipeline brings the time series from the volume into the CIFTI grayordinate standard space. This is accomplished by mapping the voxels within the cortical gray matter ribbon onto the native cortical surface, transforming them according to the surface registration onto the 32k Conte69 mesh, and mapping the set of subcortical gray matter voxels from each subcortical parcel in each individual to a standard set of voxels in each atlas parcel. The result is a standard set of grayordinates in every subject (i.e., the same number in each subject, with spatial correspondence) with 2mm average surface vertex and subcortical volume voxel spacing. These data are smoothed with surface and parcel constrained smoothing of 2mm FWHM (full width half maximum) to regularize the mapping process^61^.

### The language task

The HCP language task was developed by Binder and colleagues^34^. In the story blocks, participants were presented with brief auditory stories adapted from Aesop’s fables, followed by a 2-alternative forced-choice question to check the participants’ understanding of the story topic. The example provided in the original paper is *“For example, after a story about an eagle that saves a man who had done him a favor, participants were asked, “Was that about revenge or reciprocity?””.* In the math blocks, participants were also presented auditory series of addition and subtraction (e.g., “fourteen plus twelve”), followed by “equals” and then two choices (e.g., “twenty-nine or twenty-six”). To ensure similar level of difficulty across participants, math trials automatically adapted to the participants responses. As shown by Binder and colleagues^34^, the story and math trials were well matched in terms of duration, auditory and phonological input, and attention demand. Furthermore, they were likely to elicit distinct brain activation – on the one hand, anterior temporal lobes classically involved in semantic processing, and parietal cortex on the other hand, classically involved in numerical processing, thus spanning a broad set of regions involved in conceptual semantic processing.

### HCP task fMRI analysis

The analysis of fMRI data was carried out by the HCP consortium and we describe briefly their pipeline^13^. The Story predictor covered the variable duration of a short story, question, and response period (~30 s). The Math predictor covered the duration of a set of math questions designed to roughly match the duration of the story blocks. The grayordinate data for individual task runs were processed in a level 1 analysis. Activity estimates were computed for the preprocessed functional time series from each run using a general linear model (GLM) implemented in FSL’s FILM (FMRIB’s Improved Linear Model with autocorrelation correction)^64^. Predictors were convolved with a double gamma “canonical” hemodynamic response function^65^ to generate the main model regressors. The two runs for each task and subject were then combined in a level 2 fixed-effects analysis^13^, which we used as our phenotype. Fixed-effects analyses were conducted using FEAT (fMRI Expert Analysis Tool) to estimate the average effects across runs within-subjects, and then mixed-effects analyses treating subjects as random effects were conducted using FLAME (FMRIB’s Local Analysis of Mixed Effects) to estimate the average effects of interest for the group third-level analysis.

### Phenotype definitions

To define our phenotypes, we consider separately the regression analyses on STORY and MATH tasks, and the contrast STORY-MATH. We used the beta values (pe1.dtseries.nii files) of the results of the level 2 analysis, which essentially average the level 1, i.e., the individual, runs. The contrasts were defined by the HCP in level 1 and averaged for level 2: thus, the grayordinate values of the beta and contrast values (cope1.dtseries.nii) are identical in this case, as they did not define any “new” contrasts specifically at level 2. Therefore, we could have used the cope1.dtseries.nii.files with no difference in results. We used the MSMAll registered the functional analysis results from HCP and the HCP multimodal parcellation^15^. We analyzed each of the 180 areals separately. We computed the median beta values in each areal for both hemispheres. These phenotypes constitute our proxy to estimate the activation in each part of the brain.

Moreover, we also included in our phenotypes the accuracy, reaction time and average difficulty level for the MATH and STORY tasks. We called these “behavioral scores”, as opposed to the grayordinate activation phenotypes previously defined.

### Univariate analysis of additive genetic variance

The variance components method, as implemented in the Sequential Oligogenic Linkage Analysis Routines (SOLAR) software package^66^, was used for the heritability estimations of the phenotypes under analysis, such as the median activation in each areal^15^. The SOLAR algorithms use maximum variance decomposition methods derived from the strategy developed by Amos^67^. The covariance matrix Ω for a pedigree of individuals is given by:

Ω = 2⋅Φ⋅σ_g_² + I⋅ σ_e_², where σ_g_² is the genetic variance due to the additive genetic factors, Φ is the kinship matrix representing the pair-wise kinship coefficients among all individuals, σ_e_² is the variance due to individual-specific environmental effects, and I is the identity matrix.

Narrow sense heritability is defined as the fraction of the phenotype variance σ_p_² attributable to additive genetic factors: h² = σ_g_²/ σ_p_².

The significance of the heritability is tested by comparing the likelihood of the model in which σ_g_² is constrained to zero with that of the model in which σ_g_² is estimated. Before testing for the significance of heritability, phenotype values for each individual within the HCP cohort were adjusted for the following covariates: sex, age, age², age⋅sex interaction, age²⋅sex interaction, ethnicity (Hispanic or not) and education level. We used the number of years of education as a proxy for the education level to account for environmental differences in family socioeconomic status. This is a conservative approach because the number of years of education was shown to be associated not only with the family socioeconomic status (7%) but also with the general cognitive ability (3.5%)^68^. Thus, it likely has shared environmental ground with the former and shared genetic origin with the latter. HCP data do not contain the information that would disentangle this issue. Following this last remark, one should note that the heritability estimates and shared genetic variances, described in the next section, were underestimated.

### Bivariate genetic analyses

To assess the relationship between math dexterity/language comprehension and activation in brain areas, we computed the Pearson correlation between the median activation in each of the 180 areals of both hemispheres, and the behavioral scores.

Furthermore, we assessed the degree of shared genetic variance in the areals for which activation was significantly correlated with the behavioral scores; we performed a genetic correlation analysis using SOLAR, relying on the following model:
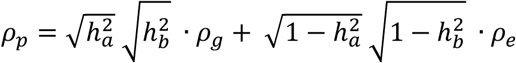 where Pearson’s phenotypic correlation ρ_p_ is decomposed into ρ_g_ and ρ_e_. ρ_g_ is the proportion of variability due to shared genetic effects and ρ_e_ that due to the environment, while 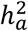 and 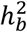 correspond to the previously defined narrow sense heritability for phenotypes *a* and *b*, respectively. In our case, one corresponds to the heritability of fMRI activation in one areal, while the second is the heritability of one of our behavioral scores.

## Acknowledgments

Data were provided by the Human Connectome Project, WU-Minn Consortium (Principal Investigators: David Van Essen and Kamil Ugurbil; 1U54MH091657) funded by the 16 NIH Institutes and Centers that support the NIH Blueprint for Neuroscience Research; and by the McDonnell Center for Systems Neuroscience at Washington University.

## Author Contributions Statement

Y.L.G and V.F designed the analysis and wrote the manuscripts. Y.L.G prepared the figures and tables. All authors analyzed the results and reviewed the manuscripts.

## Additional Information

Competing financial and non-financial interests

All the co-authors have no conflict of interest to declare.

